# Differences between Omicron SARS-CoV-2 RBD and other variants in their ability to interact with cell receptors and monoclonal antibodies

**DOI:** 10.1101/2022.01.29.478316

**Authors:** Carolina Corrêa Giron, Aatto Laaksonen, Fernando Luís Barroso da Silva

## Abstract

SARS-CoV-2 has caused immeasurable damage worldwide and available treatments with high efficacy are still scarce. With the continuous emergence of new variants of the virus, such as Omicron, Alpha, Beta, Gamma, and Delta - the so-called variants of concern, the available therapeutic and prevention strategies had to return to the experimental trial to verify their effectiveness against them. This work aims to expand the knowledge about the SARS-CoV-2 receptor-binding domain (RBD) interactions with cell receptors and monoclonal antibodies (mAbs). Special attention is given to the Omicron variant and its comparison with the others, including its sublineage BA.2 and two new ones (B.1.640.1 and B.1.640.2/IHU) recently found in France. By using constant-pH Monte Carlo simulations, the free energy of interactions between the SARS-CoV-2 receptor-binding domain (RBD) from different variants and several partners (Angiotensin-Converting Enzyme-2 (ACE2) polymorphisms and several mAbs) were calculated. It was evaluated both the impact of mutations for the RBD-ACE2 and how strongly each of mAb can bind to the virus RBD, which can indicate their therapeutic potential for neutralization. RBD-ACE2-binding affinities were higher for two ACE2 polymorphisms typically found in Europeans (rs142984500 and rs4646116), indicating that these types of polymorphisms may be related to genetic susceptibility to COVID-19. The antibody landscape was computationally investigated with the largest set of mAbs so far in the literature. From the 33 studied binders, groups of mAbs were identified with weak (e.g. S110 and Ab3b4), medium (e.g. CR3022), and strong binding affinities (e.g. P01’’’, S2K146 and S230). All the mAbs with strong binding capacity could also bind to the RBD from SARS-CoV-1, SARS-CoV-2 wt, and all studied variants. These mAbs and especially their combination are amenable to experimentation and clinical trials because of their high binding affinities and neutralization potential for current known virus mutations and a universal coronavirus.

## 1. Introduction

After more than two years of struggling against COVID-19 and all the sense of loss caused by it (not only because of the number of deaths but also because of all the plans that had to be canceled or postponed, the mental struggle, economic and pedagogical impacts, long-term effects, etc), the new SARS-CoV-2 variants have been continuously reignited a security alert and brought doubts about the efficacy of the therapeutical and prevention strategies already developed (Iyer, 2021). At the beginning of the pandemic, in Manaus, Brazil, even though studies indicated that 76% of the population had already been infected before the outbreak of the Gamma variant (P.1), in the so-called “first wave”, there was a resurgence of COVID-19 in the city despite high seroprevalence (Sabino et al., 2021). All these emerging variants, especially the ones called VOC (variants of concern) by some international organizations, such as the World Health Organization (WHO) (Parums, 2021; *Tracking SARS-CoV-2 Variants*, n.d.) - Alpha, Beta, Gamma, and Delta (as of September 2021) - are threatening because of the impact the mutations can have on binding, entry and replication mechanisms, resulting in possible immune escapes, making reinfections possible and demanding the already developed treatments to be updated (Kupferschmidt, 2021a; Plante et al., 2021). However, SARS-CoV-2 is still a newly discovered virus, thus it is still adapting to its human host, requiring close attention to mechanisms that it may develop (Plante et al., 2021).

A recent threat is being caused by the new variant Omicron (B.1.1.529), a highly divergent variant with a high number of mutations and designated VOC by WHO first identified in South Africa at the end of November 2021 and responsible now by daily records of new cases on a global scale (Kupferschmidt, 2021b). Omicron, the fifth VOC, apparently did not develop out of the previous VOC due to its genomic difference (CDC COVID-19 Response Team, 2021; Kupferschmidt, 2021b; O’Toole et al., 2021; World Health Organization, 2021). Among other possibilities, likely, it evolved early somewhere with insufficient infrastructure to test and supervise (Kupferschmidt, 2021b). Omicron has a different set of mutations in the Spike protein, containing the substitution of 30 amino acids, insertion of three residues, and deletion of six residues (Dejnirattisai, Huo, et al., 2021; Dejnirattisai, Shaw, et al., 2021), some of which may be associated with much higher transmissibility and humoral immune escape potential (Dejnirattisai, Huo, et al., 2021). The viral load does not seem superior to what was seen before for Delta (Kozlov, 2021).

Despite the newness of the variant, scientists are racing to understand the worrisome threat it poses and also the impact of its sublineage BA.2 (not yet classified as a VOC) (Metzger et al., 2021). Some initial studies suggested that the mutations found in Omicron might affect the antibodies’ neutralization ability (Dejnirattisai, Shaw, et al., 2021) although it is not clear how immune responses triggered by either a prior infection or vaccination could still offer some protection (Callaway & Ledford, 2021). At the moment, due to the contrast to the original SARS-CoV-2 sequence, it seems that the vaccines produced with the original strain are less effective (Cele, Jackson, et al., 2021; Dejnirattisai, Shaw, et al., 2021; Dejnirattisai, Huo, et al., 2021; Rössler et al., 2021). The same can be seen for the plasma neutralization of the SARS-CoV-2 Omicron variant using blood from patients previously infected with other variants (Schmidt et al., 2021). Inactivated-virus vaccines, such as those made by Sinovac and Sinopharm of China, Covaxin of India, COVIran Barekat of Iran, and QazVac of Kazakhstan, were shown to provide little or no protection against infection with the Omicron variant (Dolgin, 2022; Lu et al., 2021). Even two jabs fail to produce antibodies (Abs) against the VOC (Dolgin, 2022). The third dose of another type of vaccine, such as those based on messenger RNA or purified proteins, appears to offer better protection (Dolgin, 2022; Pérez-Then et al., 2021; Zuo et al., 2022). However, there is still no evidence of increased severity with Omicron (Dejnirattisai, Shaw, et al., 2021).

Preliminary results from different labs indicate a decreased lung cell infectivity in animal models, suggesting that Omicron might be less virulent than its ancestral (Abdelnabi et al., 2021; Bentley et al., 2021; Diamond et al., 2021; Meng et al., 2021). Initial clinical data could not yet access if Omicron is less pathogenic or is mild virulence due to preexisting acquired or natural immunity (Maslo et al., 2021). Nevertheless, the health risk remains critical due to its extraordinarily high transmissibility, which can simultaneously bring a large number of people to the hospitals, overloading the health systems. It can also result in more deaths despite its lower mortality rate because more people can be contaminated. It remains to be seen whether available monoclonal antibodies (mAbs) either vaccine-induced or from convalescent patients or industrially produced can neutralize Omicron. Initial data are not promising as several papers and preprints show escapes for the majority of existing SARS-CoV-2 neutralizing Abs (Cao et al., 2021; Dejnirattisai et al., 2022; Kozlov, 2021; Sokal et al., 2021).

Several independent factors are known to increase viral transmission and the development of the most severe form of the disease, whether they are related to an increased binding affinity of the virus to specific cell receptors or higher stability of the molecular protein complexes responsible for some specific task within the virus life cycle. Some of them are related to the virus itself and its mutations, others to external factors, and some are related to the individual. Among the factors related to the individual, it can be mentioned the genetic ones, which include **i**) the amount in the body of the specific cell receptor used by SARS-CoV-2 to enter the human cells [i.e. the Angiotensin-Converting Enzyme-2 (ACE2) receptors explaining, for instance, the higher prevalence of the disease in men (Packham, 2020)], **ii**) the role that ACE2 expression plays in the host range and the tropism of SARS-CoV-2 in human tissues (Gao & Zhang, 2020), and **iii**) the frequency of human leukocyte antigen (HLA) genotypes - such as HLA-A*11:01 - which is associated with the fatality rates (Toyoshima et al., 2020). From the virus perspective, some SARS-CoV-2 variants are known to have mutations that energetically favor the RBD-ACE2 complexation, which could increase infectivity (W. Hu et al., 2021; A. Khan et al., 2021). Others can favor an entry to infect the human cell by a stronger affinity of their spike protein with the host protease furin (Whittaker, 2021). The number of spike proteins at the “open state” can also play an important role (Giron et al., 2021). The balance between these entering mechanisms might define how much a given variant will be infectious and transmissible.

Regarding external factors, there are social human behaviors, such as disrespecting social isolation and not wearing personal protective masks that increase the transmissibility, vaccine hesitancy, slow vaccinate rates, etc., but specific mutations in the viral proteins can also do so, and some mutations are even related to immune system evasion. Some SARS-CoV-2 variants show convergent mutations. The Beta (South African variant/B.1.351) and the Gamma (Manaus variant/P.1) carry the same key mutations in the spike protein, even though they emerged from independent events: **a**) N501Y, also present in the Alpha (UK/B.1.1.7) and Omicron (B.1.1.529) variants, and **b**) K417T/K417N, and E484K, that are known to play a role in the immune escape (Wibmer et al., 2021). E484K has a high effect on the interaction with mAs and human sera, reducing neutralization capacity (Greaney et al., 2021; Uriu et al., 2021; Voloch et al., 2021; Weisblum et al., 2020). N501Y, on the other hand, increases transmissibility because of its enhanced binding affinity with the cellular receptor ACE2, not having large effects on antibodies’ neutralization alone (Starr et al., 2020). However, combined with K417T and E484K, there is a synergetic effect and a substantially decreased neutralization of mAbs as verified in experiments, which is also greater than the effect produced by each mutation separately (Cele, Gazy, et al., 2021; Z. Wang et al., 2021; Wibmer et al., 2021). In Omicron, these mutations appear as K417N (as seen in Gamma) and E484A together with N501Y and other 12 replacements at the RBD. There is a frequent tendency that mutations involve the replacement of the physical-chemical nature of acid and basic residues.

It is difficult to control the external factors in many countries and impossible to modify individual genetic characteristics. Together with the vaccines (Mallapaty et al., 2021), therapeutical interventions are the most effective means to effectively and practically reduce the number of deaths (Hastie et al., 2021; Reis et al., 2022; Tso et al., 2021). mAbs offer a promising route for clinical treatments under a booming expansion (J. Chen et al., 2021; Marovich et al., 2020; Mullard, 2021). All these SARS-CoV-2 variants require continuous analysis of the available mAbs and often the development of new and specific ones. In addition, it is necessary to carry on molecular studies of the biomolecular interactions related to the virus’ proteins to understand their relationship with the transmission, infectivity, virulence, lethality and also to see other possible intervention routes. Due to the similarity between SARS-CoV-2 and SARS-CoV-1 spike proteins (identity and the similarity are 64.8% and 78.0%, respectively, as obtained by the alignments at the server EMBOSS Needle (Needleman & Wunsch, 1970) with default settings) and the similar entry mechanism (via ACE2 receptor) (Dimitrov, 2003; Walls et al., 2020), the first mAbs tested for the COVID19 virus were the ones that had already been known for the 2003 Severe Acute Respiratory Disease outbreak (SARS-CoV-1 virus), such as 80R and S230 (Du et al., 2009; Giron et al., 2020; Needleman & Wunsch, 1970; Shang et al., 2020; X. Tian et al., 2020). S230, for example, was found to block SARS-CoV-1 binding to the ACE2 receptor with a high affinity although it seems to mimic ACE2 so well that it can trigger fusogenic conformational changes through functional receptor mimicry (Walls et al., 2019). Yet, it is not clear whether S230 would be able to block and eventually neutralize Omicron without such undesired negative effects that could favor the virus break into cells, and consequently, it can be severely harmful to the patients. Cao and co-authors reported in a preprint that most of the available mAbs showed reduced efficacy (over 85% out of the 247 human anti-RBD tested binders are escaped by Omicron) (Cao et al., 2021). S230 was not yet analyzed in their high-throughput yeast display screening and pseudovirus neutralization assays. Promising therapeutic binders found in Caós study against Omicron were S309 and CR3022. Theoretical studies also suggested that CR3022, the engineered CR3022-like mAb, P1, and P1’’’ seem to be good candidates for Omicron (Neamtu et al., 2022). Another experimental group has shown that S309 was minimally affected by the multitude of mutations found in Omicron (VanBlargan et al., 2021). Conversely, assays with Vero-TMPRSS2 and Vero-hACE2-TMPRSS2 cells have shown that several other mAbs (including LY-CoV555, LY-CoV016, REGN10933, REGN10987, and CT-P59) completely lost their inhibitory activities for Omicron (VanBlargan et al., 2021). It is also not completely clear if all molecules that are powerful in the binding are also so efficient in virus neutralization (Chi et al., 2020). Yuan and collaborators pointed out the urgency for the next generation of vaccines and therapeutic medicines effective against Omicron after comparing its pathogenesis with the Delta variant in the golden Syrian hamster COVID-19 model (Yuan et al., 2022). The good news is that the cross-reactivity seems possible, specifically between Abs for Delta and Omicron variants (K. Khan et al., 2021).

Once the virus can potentially develop resistance to a single antibody due to an accumulation of mutations in some epitope regions (Renn et al., 2020), cocktails are an interesting therapeutic option. There are some cocktails commercially developed, such as REGN-COV2, which consists of two mAbs that bind to different regions of the Spike (S) protein (Baum et al., 2020). REGN-COV2 provided benefits in trials with rhesus macaques and golden hamsters, simulating mild and severe disease, respectively, enhancing viral clearance, and was released by some national health agencies for emergency use, such as Anvisa, the Brazilian agency (Baum et al., 2020; Weinreich et al., 2021). LY-CoV555, another promising neutralizing antibody that entered clinical evaluation, was also studied in a cocktail form with LY-CoV016 (P. Chen et al., 2021; Renn et al., 2020; Starr et al., 2021). However, it was found that the mutation E484K (which is present in B.1.351 and P.1 SARS-CoV-2 variants) and L452R (found in the B.1.429 variant) escape LY-CoV555, and the mutations K417N/T escape LY-CoV016 (Starr et al., 2021). Moreover, individual mutations were found to escape the cocktail LY-CoV555+LY-CoV016 (Starr et al., 2021). Apparently, Omicron can also evade them (Cao et al., 2021).

Due to the emergence of these new variants, especially the highly-transmissible Omicron variant, efforts are being concentrated to understand the impact of the mutations in different steps of the virus cycle, and the consequent transmission, virulence, and infectivity. Here, we extended our previous *in silico* study (Giron et al., 2020) to investigate and compare the impact of these mutations on the interaction between the RBD and the human ACE2 since this is the very first step for the SARS-CoV-2 cellular entry. ACE2 polymorphisms were also included in our analysis as they were suggested to be a genetic factor that could affect the viral entry independently, but also the severity of the disease due to its role in comorbidities, such as hypertension (F. Chen et al., 2021; Möhlendick et al., 2021). ACE2 rs2285666, for example, is associated with increased infection risk and fatality (Möhlendick et al., 2021). Another study suggests that T92I and K26R ACE2 mutations favor the binding with SARS-CoV-2, whereas K31R, E37K, and others decrease the binding affinity (Suryamohan et al., 2021). Furthermore, hypomethylation and zinc deficiency, common amongst the elderly, can enhance the expression of ACE2, which can help explain why they are more susceptible to developing the most severe form of the disease (F. Chen et al., 2021).

Several recently discovered mAbs whose structure has been made available in the literature were also investigated (LY-CoV555/Bamlanivimab, LY-CoV016/Etesevimab, 4A8/PGC-1α, SARS2-38, AZD1061/Cilgavimab, AZD8895/Tixagevimab, chAb25, COV2-2196, S309 (the parent mAb of Sotrovimab (McCallum et al., 2021)), S230, REGN10933/Casirivimab, REGN10987/Imdevimab, CT-P59, EY6A, COR101, Fab15033, B38, m336, and S110) for these variants. The main aim of the present work is to compare Omicron with other strains in terms of how its RBD binding affinities are for both the complexation with ACE2 polymorphisms and the mAbs. Similarities and differences between mutations in these molecules and the antibody landscape (Hachim et al., 2021) were quantified which allowed indicating the possible efficacy of these mAbs can be against the VOCs, other variants of interests (VOI) (Parums, 2021), and additional specific residue replacements. Such information sheds light on the physiopathology of COVID19 and on future directions to be explored for efficient antibody products for clinical use.

## 2. Materials and Methods

*In silico* methods have proven their value in different research fields, including virology, immunology, and biomolecular engineering (Batra et al., 2020; Ibrahim et al., 2018; Lousa et al., 2022; Sato et al., 2013; Sharma et al., 2015). Among the available tools (Barroso da Silva et al., 2020; Germain et al., 2011; Schlick & Portillo-Ledesma, 2021), Molecular Dynamics and Monte Carlo (MC) simulations can be widely used to explore the multiple aspects of virus infection and disease, such as pathogenesis, immunology, the stability of the molecular structures, as well as their intermolecular interactions.

### a. Molecular simulations

Here, a robust and simplified theoretical methodology designed to explore protein-protein interactions at constant-pH conditions is used to investigate the biomolecular interactions between both the host and pathogen and the antigen-antibody. This coarse-grained (CG) model combines translational and rotational motions of two rigid macromolecules in a cylindrical simulation box with the “Fast Proton Titration Scheme” (FPTS) (Barroso da Silva & MacKernan, 2017). Solving this CG model in an MC simulation, the free energy of interactions for the complexation can be estimated for different pairs of macromolecules at different physical-chemical conditions at lower CPU costs in comparison with other models (Neamtu et al., 2022). Two pairs of macromolecules were investigated: **a**) RBD from different variants with ACE2 with sequences from key polymorphisms (RBD_x_-ACE2_x_ where *x* is either the wildtype [wt] or a mutated form), and **b**) RBD from different variants with several known mAbs (RBD_x_-mAb_y_ where *y* is one of the 33 studied binders -see the list below). Therefore, an extensively large number of simulations have to be performed which would be impossible with the inclusion of additional degrees of freedom in the model. The transition between the “up/down” states of the spike protein is not incorporated in the simulated model. It is assumed that the spike protein was at the “open” configuration, when it is highly accessible to ACE2 and/or the other binders (Giron et al., 2020). Thinking about the suggested two-step “expose–dock-like” mechanism (Giron et al., 2020, 2021), we focused here on the second step, after conformational adjustments were completed by the spike homotrimer to allow the complexation RBD-ACE2 to happen with no steric clashes.

pH is an input parameter and the charges of all titratable groups can be changed due to the protonation-deprotonation process during the simulation runs. This implies that the electrostatic interactions are properly taken into account in the model including mesoscopic physical mechanisms that can be responsible for an additional attraction (Barroso da Silva et al., 2006; Kirkwood & Shumaker, 1952). This methodology has been successfully applied in several biomolecular systems from food proteins to septins (Barroso da Silva et al., 2018; Delboni & Barroso da Silva, 2016; Mendonça et al., 2019; Persson et al., 2010). It has also been used to investigate the flaviviruses (Poveda-Cuevas et al., 2018, 2020) and coronavirus (Giron et al., 2020, 2021; Neamtu et al., 2022). Recently, this theoretical approach was named FORTE (Fast cOarse-grained pRotein-proTein modEl) (Neamtu et al., 2022).

As discussed before (Poveda-Cuevas et al., 2018, 2020), intrinsic assumptions are **a**) electrostatic properties are of great importance in promoting interactions between macromolecules (these aspects seem to be even more critical for virus proteins (Bozek et al., 2012; Giron et al., 2020, 2021; Ishikawa et al., 2021; Nanda et al., 2010; Neamtu et al., 2022; Poveda-Cuevas et al., 2018, 2020; Ramaraj et al., 2012), **b**) the molecular sequence-structure relationship is powerful enough to introduce mutations or modifications while preserving the overall coiling of these molecules (structural comparisons between the wt and mutated ones confirmed that this is not a critical assumption especially for the spike protein and the RBD - see (Giron et al., 2021; Neamtu et al., 2022), and **c**) conformational macromolecular changes have a minor impact in their binding free energies in particular for medium and long-range separation distances (Neamtu et al., 2022; Wade et al., 1998). This last item is less critical as the closer the physical-chemical conditions of the simulations are from the used experimental coordinates. It is worth mentioning that the FORTE approach provides derivatives of the free energy (e.g., the interaction free energy as a function of separation distances) and are not limited to a specific separation/orientation between the macromolecules nor exclusively in crystallographic experimental conditions. This is different and far more comprehensive than classical “docking” procedures, where the focus is on short-range events (Giron et al., 2020; Neamtu et al., 2022). The reader is referred to these references for more information.

Briefly, the model considers each amino acid as a single charged Lennard-Jones (LJ) sphere of radius (*R_i_*) with valence *z_i_*, and initial coordinates as given by its three-dimensional structures (see below). The values of *R_i_* for each type of amino acid were taken from Ref. Persson et al. (2010). Such different sizes of the residues partially account for the non-specific contributions from the hydrophobic effect in the model (Delboni & Barroso da Silva, 2016; Poveda-Cuevas et al., 2020). This is important to preserve the macromolecular hydrophobic moments (Eisenberg et al., 1982) and contribute to guiding a correct docking orientation at short separation distances. All valences of titratable groups z_i_ are pH-dependent and controlled by the FPTS (Barroso da Silva & MacKernan, 2017; Barroso daSilva & Dias, 2017; Teixeira et al., 2010). These valences can vary during the simulation. Non-titratable amino acids are assigned with a zero charge kept constant all the time. Two ionized amino acids of valences *z_i_* and *z_j_* (either found in the same chain or in the other macromolecule) contribute to the electrostatic potential energy as given by,

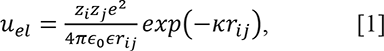

where *r_ij_* is the inter-residues separation distance, *κ* is the modified inverse Debye length (Barroso da Silva & MacKernan, 2017; Jurado de Carvalho et al., 2006; Teixeira et al., 2010), *ε_s_* is the dielectric constant of the medium (78.7 to mimic an aqueous solution at temperature T=298K), *ε_0_* is the dielectric constant of the vacuum (ε_0_=8.854×10^− 12^ C^2^/Nm^2^), and *e* is the elementary charge (*e*=1.602×10^− 19^C). Further details can be found in Refs. Barroso da Silva et al. (2016); Delboni and Barroso da Silva (2016); Barroso da Silva et al. (2018); Poveda-Cuevas et al. (2020). As mentioned before, the charges for the ionisable amino acids were defined by the FPTS method (Barroso da Silva & MacKernan, 2017; Teixeira et al., 2010).

Together with electrostatic interactions, the simulation model also includes the effects of other physical contributions, such as hydrophobic effect, van der Waals interactions, and excluded volume repulsion, through an LJ term [u_vdw_(r_ij_)] between the residues (Barroso da Silva et al., 2016; Delboni & Barroso da Silva, 2016; Poveda-Cuevas et al., 2020):

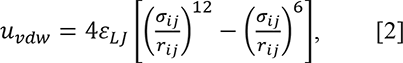

where *ε_LJ_* regulates the attractive forces in the system (Barroso da Silva et al., 2016; Delboni & Barroso da Silva, 2016; Poveda-Cuevas et al., 2020) and was assigned the universal value of 0.124 kJ/mol (Barroso da Silva et al., 2016; Delboni & Barroso da Silva, 2016; Hyltegren et al., 2020; Persson et al., 2010), and *σ_ij_* is given by Lorentz-Berthelot rule from *R_i_* and *R_j_* (Leach, 2001).

The full configurational energy of the system then becomes

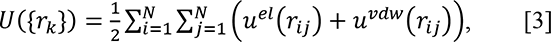

where {r_k_} are amino acid coordinates at a given configuration and *N* is their total number. This model was stochastically solved by Metropolis MC simulations with physiological ionic strength (150 mM) and pH 7 at 298 K using a simulation cylindrical cell of the radius of 150Å and height equals 200 Å. Some calculations were also run at pH 4.6 to investigate the pH effect for selected systems. At least 3.10^9^ MC steps were performed during the production phase. During this phase, the histogram *hist(r)* was built, which contains the number of times the two macromolecules were observed at a separation distance between *r* and r+dr (*dr* is the histogram bin size). A value of 1Å was used for *dr*. Such histogram was converted to a pair radial distribution function [g(r)] from where the free energy of interactions for the complexation process [ꞵw(r), where ꞵ is equal to the reciprocal of K_B_T and *K_B_* is the Boltzmann constant] as a function of *r* was calculated (Allen & Tildesley, 1989; Tuckerman, 2010). Even if FORTE contains some simplifications to decrease the CPU time, the need to populate all the histogram bins used for the g(r) during the sampling, the reduction of the statistical noises in the calculation of ꞵw(r), and the free energy barriers of the systems turn to be computationally demanding (Barroso da Silva et al., 2016; Delboni & Barroso da Silva, 2016; Poveda-Cuevas et al., 2020). Three replicates per simulated system were used in order to estimate the standard deviations.

### b. Molecular systems and their structural modeling

This work investigated numerous molecular systems, such as SARS-CoV-2 S RBD proteins with ACE2 (wild-type and with single/double mutations due to its most common ACE2 polymorphism: D355N, E37K, K26R, Y50F, D38V, G326E, Y83H, D509Y, H34R, K31R, E23K, H378R, K68E, E35K, I31K, N64K, and K26R+I31K) and fragments of some mAbs (see the list below). The investigated ACE2 variants here were chosen on the basis of the work of Suryamohan and co-authors (Suryamohan et al., 2021) that analyzed a large number of samples from public genomic datasets with a broad representation of the population. Our study was limited to ACE2 variants involving only the replacement of ionizable amino acids due to the nature of our CG model. The three-dimensional structures of most of these macromolecules (RBDs, ACE2, and mAbs) were obtained from the RCSB Protein Data Bank (PDB) (Berman, 2000), and all non-terminal missing regions in these proteins were built up using before calculations by the tool “Modeller” with default parameters (Eswar et al., 2006) from the program “UCSF Chimera 1.14” (Pettersen et al., 2004). All the heteroatoms and water molecules were removed from the PDB files. The GLYCAM PDB pre-processor webtool (Grant et al., 2020) was used to assign the cysteines involved in disulfide bonds that cannot titrate and to perform basic checks. The coordinates for ACE2 (wt), SARS-CoV-1 RBD, and the fragments of mAbs - LY-CoV555, LY-CoV016, LY-CoV1404, 4A8, SARS2-38, AZD1061, AZD8895/COV2-2196, chAb25, S309, S230, REGN10933, REGN10987, CT-P59, EY6A, COR-101, Fab15033, B38, m336, m396, F26G19, CR3022, P2B-2F6, 80R, BD23, H11D4, and S2K146 were obtained from PDB ids 7KMG (chains A and B, pH 4.6), 7C01 (chains H and L, pH 7), 7MMO (chains A and B, pH 6.0-7.0), 7C2L (chains H and L, pH 8), 7MKM (chains H and L, pH 7.4), 7L7E (chains M and N, pH 4.2), 7L7D (chains H and L, pH 8.0-8.5), 7EJ4 (chains H and L, pH 7.6), 6WS6 (chains A and B, pH 8), 6NB6 (chains H and L, pH 8), 6XDG (chains B and C, pH 7.4), 6XDG (chains D and E, pH 7.4), 7CM4 (chains H and L, pH 8), 6ZER (chains B and G, pH 8), 7B3O (chains H and L, pH 5.6) (Bertoglio et al., 2021), 7KLH (chains I and M, pH 7.2), 7BZ5 (chains H and L, pH 6), 4XAK (chains H and L, pH 7.5), SDD8 (chains H and L, pH 6.5), 3BGF (chains B and L, pH 5.5), 6W41 (chains H and L, pH 4.6), 7BWJ (chains H and L, pH 4), 2GHW (chain B, pH 4.6), 7BYR (chains H and L, pH 7) (Cao et al., 2020), 6Z43(chain D, pH 8.0), 7TAT (chains D and E, pH 8) respectively. These fragments of mAbs are going to be referred to simply as mAbs from now on. For a proper comparison of the outcomes, the SARS-CoV-2 RBD (wt) coordinates were taken from previous works (Giron et al., 2020, 2021) - also available at Zenodo (*Build Structural Models for Several Variants of SARS-CoV-2 RBD | Zenodo*, n.d.). Some variable new antigen receptors (VNARs) were also tested: C02, and 3B4, obtained from PDB id 7SPP (chain B, pH 7.4), and 7SP0 (pH 7.4), respectively (Ubah et al., 2021). S110 was constructed at the interactive workspace of the SWISS-MODEL workspace (Waterhouse et al., 2018). CR3022’, P01, and P01’’’ are engineered mAbs taken from previous works (Giron et al., 2020; Neamtu et al., 2022). Figure 1 shows their pair sequences alignment as obtained by the server EMBOSS Needle (Needleman and Wunsch, 1970) with default settings.

**Figure 1:**
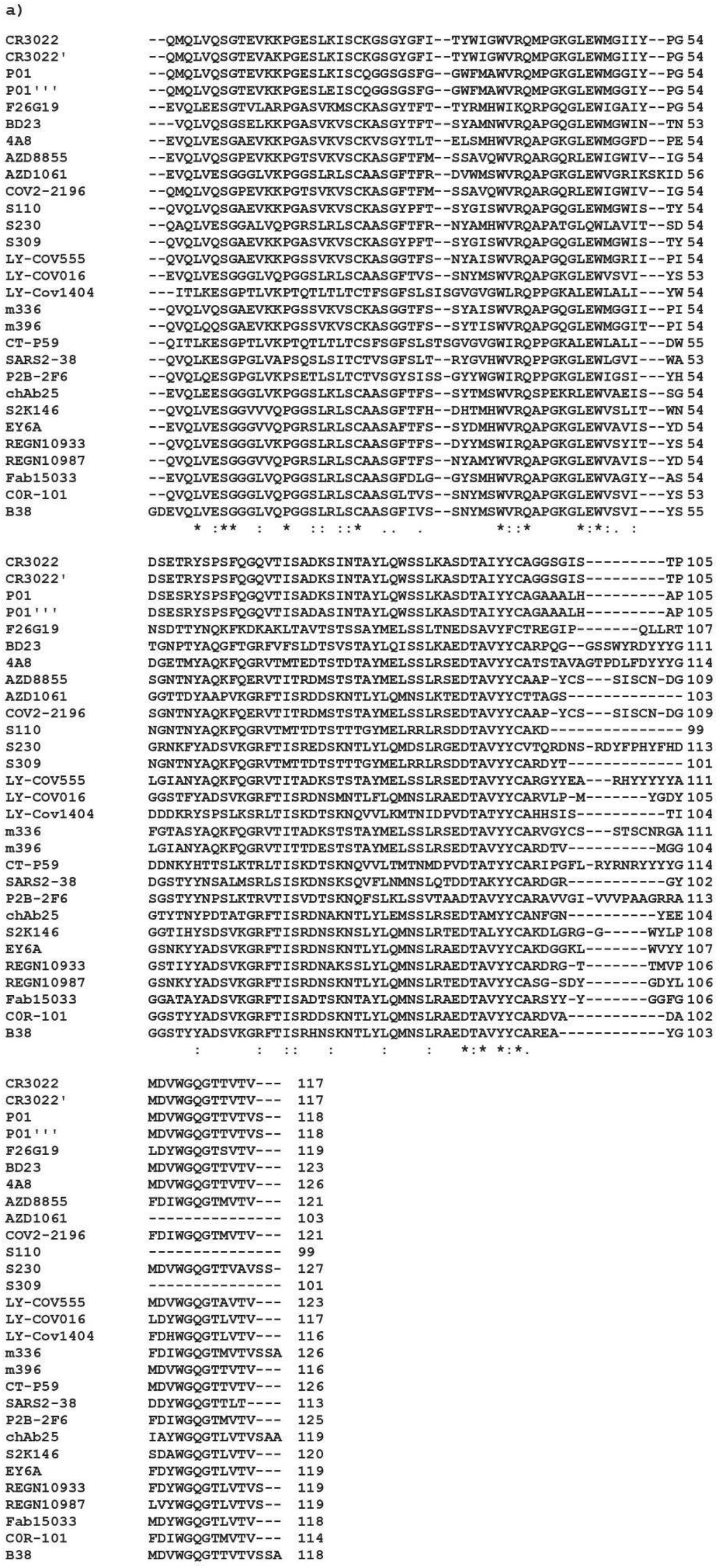
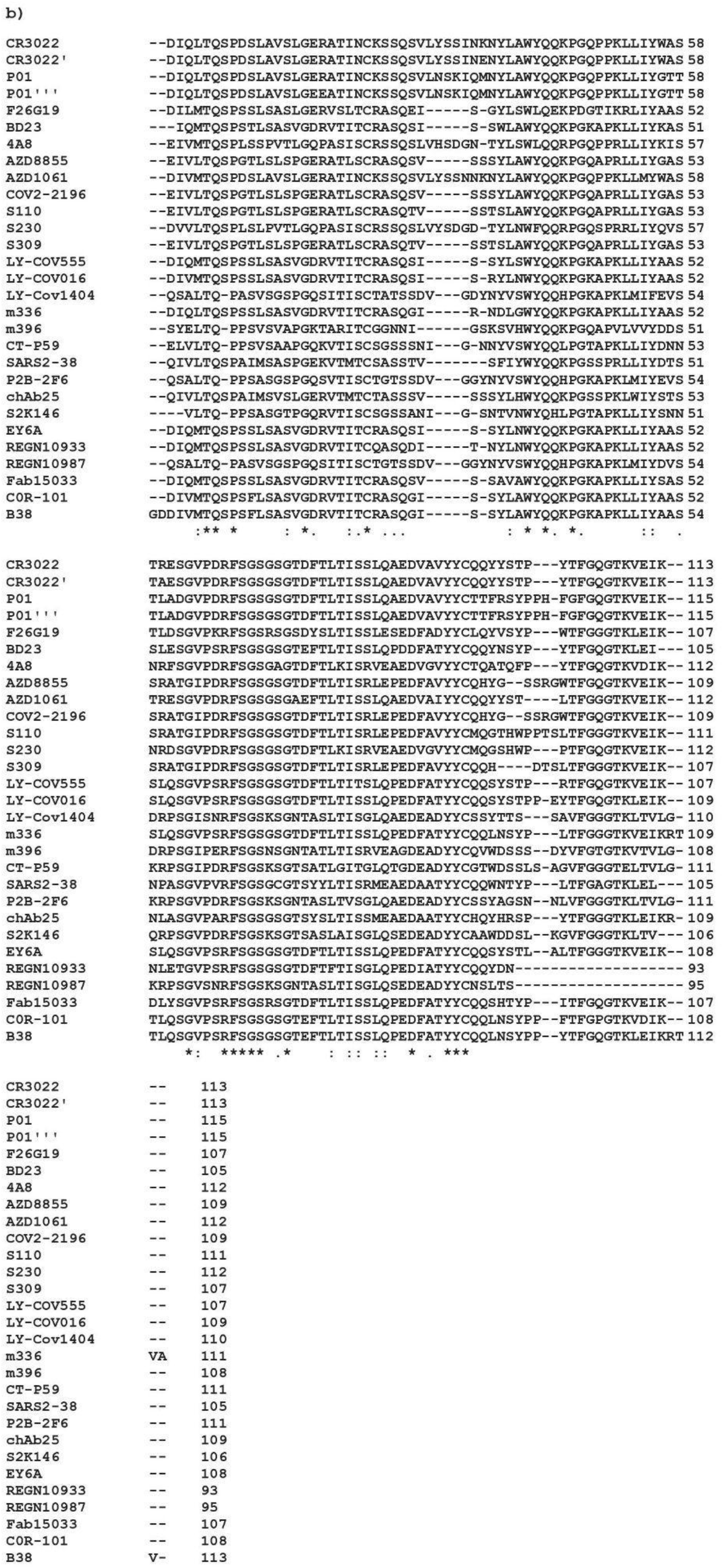
Sequences of the a) Heavy chains and b) Light chains of the studied mAbs. Dots represent identities. Symbols between aligned sequences have the usual meaning.

**Table 1.** Mutations in the RBD of the spike protein present on each studied variant. Titratable residues are highlighted in bold.

| *Variant* | *Mutations in the RBD of the spike protein* |
| --- | --- |
| Alpha (B.1.1.7/UK) | N501**Y** (T.-J. Yang et al., 2021) |
| Beta (B.1.351/SA) | **K**417N, **E**484**K**, and N501**Y** (Tegally et al., 2021) |
| Gamma (P.1/BR) | **K**417**T**, **E**484**K**, and N501**Y** (Voloch et al., 2021) |
| Delta (B.1.617.2/India) | L452**R**, T478**K**, and **E**484**Q** (J. Zhang et al., 2021) |
| Omicron (B.1.1.529/SA) | G339**D**, S371L, S373P, S375F, **K**417N, N440**K**, G446S, S477N, **T**478**K**, **E**484A, Q493**R**, G496S, Q498**R**, N501**Y**, and **Y**505**H** (Ford et al., 2021) |
| Epsilon (B.1.427/B.1.429) | L452**R** (W. Zhang et al., 2021) |
| Eta (B.1.525) | **E**484**K** (Pereira et al., 2021) |
| Iota (B.1.526) | **E**484**K**, and N501**Y** (West et al., 2021) |
| Kappa (B.1.617.1) | L452**R**, and **E**484**Q** (J. Zhang et al., 2021) |
| Mu (B.1.621) | **R**346**K**, **E**484**K**, and N501**Y** (X. Xie et al., 2021) |
| B.1.640.1 | **R**346S, N394S, **Y**449N, F490**R**, and N501**Y** (Colson et al., 2021) |
| B.1.640.2 (IHU) | **R**346S, N394S, **Y**449N, **E**484**K**, F490S, and N501**Y** (Colson et al., 2021) |
| BA2 (Omicron sublineage) | S371F, T376A, **D**405N, and **R**408S (Metzger et al., 2021) |
| C1.2 | **Y**449**H**, **E**484**K**, and N501**Y** (Scheepers et al., 2021) |
| mink | **Y**453F (Bayarri-Olmos et al., 2021) |

## 3. Results and Discussion

### 3.1 Binding affinities for RBD from different variants with ACE2_wt_

The interaction with a cell receptor is typically the very preliminary step necessary for a virus to gain access to a human cell. It is consequently an important virulence trigger. At the beginning of the COVID19 pandemic, it was natural to hypothesize that SARS-COV-2 would utilize a similar cell entry mechanism via ACE2, as done by SARS-CoV-1 (Dimitrov, 2003; Walls et al., 2020). Different experimental assays lately confirmed it (X. Tian et al., 2020; Walls et al., 2020). In pioneering theoretical work at that time, Giron and co-authors investigated the interactions between the RBD (wt/Wuhan sequence) and ACE2_wt_ employing FORTE and using a molecular model built up at the SWISS-MODEL workspace (YP_009724390.1) based on the NCBI reference sequence NC_045512 (Giron et al., 2020). The experimental binding behavior was reproduced, the electrostatic epitopes (Poveda-Cuevas et al., 2020) were mapped both for SARS-CoV-1 and 2 (wt), and it became clear the importance of the electrostatic interactions for this process (Giron et al., 2020). Later, the binding and the key contribution from the electrostatic interactions were confirmed by other research groups too using additional theoretical approaches (Bai & Warshel, 2020; K. Khan et al., 2021; Morton & Phillips, 2020; D. Wang et al., 2021; Y. Xie et al., 2020, 2021). After the surge of the new variants, RBDs with the corresponding mutated sequences started to be investigated both theoretically and experimentally (Abdool Karim & de Oliveira, 2021; Davies et al., 2021; Hoffmann, Arora, et al., 2021; Starr et al., 2020; Tao et al., 2021; Toyoshima et al., 2020). In a landmark experimental research work, Starr and co-authors performed a deep mutational scanning of SARS-CoV-2 RBD quantifying the effect of every single mutation on the binding, stability, and expression (Starr et al., 2020). Yet, synergistic or antagonistic effects due to more than one simultaneous mutation were not explored. Subsequently, this was partially done by Barton and colleagues for Alpha, Beta, and Gamma VOCs (Barton et al., 2021). In parallel with the experimental works, a large number of computational studies have been conducted to investigate the effects of both individual amino acid substitutions or more simultaneous replacements (Fratev, 2021; Giron et al., 2021; Jafary et al., 2021; A. Khan et al., 2021; Lan et al., 2022; Maher et al., 2022; F. Tian et al., 2021). Recent manuscripts deposited or published during the writing of this article already reported data for Omicron (Ford et al., 2021; Han et al., 2022; Hanai, 2022; Lan et al., 2022; Lupala et al., 2022; Nie et al., 2022; Ortega et al., 2021).

Here, free energies of interactions between the RBD of important variants (both cases studied before by different methods and others yet not fully explored in the literature) with ACE2_wt_ were calculated by the FORTE approach (see Figure 2). To our knowledge, no previous studies have investigated and compared the RBD-ACE2 binding affinity for such a large number of cases under the same conditions and using a common *in silico* strategy for all of them at the same time. In Figure 2a, *ꞵw(r)* as calculated by FORTE is given for all studied RBDs and ACE_wt_ at pH 7 and physiological salt concentration. All VOCs and VOIs RBDs have a higher affinity for the ACE2_wt_ in comparison with the RBD wildtype. This comparison is easier seen in Figure 2b, where the minima free energy values (*ꞵw_min_*) found in the estimated binding affinities are plotted as bars for them. Evolution has improved the viral fitness of SARS-CoV-2 to bind to ACE2_wt_ over time. From Alpha to Omicron (data shown in blue), the affinity has increased in comparison with the wt (data shown in purple). This behavior is similar to what was shown for Alpha, Beta, Gamma, Eta, and Iota variants by Khan and co-authors in their computation work using Haddock (A. Khan et al., 2021). The values of *ꞵw_min_* for these VOCs and VOIs are also higher than what was measured for SARS-CoV-1 RBD (data shown in red). The two variants (B.1.640.1 and B.1.640.2/IHU) that recently (December 2021) were found in France (Colson et al., 2021) do not indicate a more worrying scenario since their RBDs behave similarly to what was seen by the previous VOCs. Within the statistical uncertainties, the binding affinities displayed in Figure 2b can be grouped from the lowest to the highest as group 1 (WT, B.1.640.1, B.1.640.2, BA.2, mink, and Alpha) < group 2 (SARS-CoV-1, Beta, Epsilon, Gamma, and Mu) < group 3 (Eta, Kappa, Iota, Mu-K417T, C1.2, and L452R+T478K) < group 4 (Delta) < group 5 (Omicron). The RBD_Omicron_ is clearly the system with the highest affinity for ACE2_wt_. This result for Omicron is qualitatively in agreement with several other theoretical calculations that are appearing in the literature (Ford et al., 2021; Han et al., 2022; Hanai, 2022; Lan et al., 2022; Lupala et al., 2022; Nie et al., 2022, 2022; Ortega et al., 2021). For example, Ortega and colleagues reported the values of − 109.8(3.5) and − 163.8(4.1) for the Haddock score function for the wt and Omicron, respectively (Ortega et al., 2021). Hanai predicted the following order for some variants (in bold cases that appear in the same order with the present calculations are highlighted): **Alpha** (276.9 kcal mol^− 1^) < **Beta** (289.2 kcal mol^− 1^) = **Gamma** (300.1 kcal mol^− 1^) < WT (301.9 kcal mol^− 1^) < Mu (406.8 kcal mol^− 1^) < **Kappa** (503.5 kcal mol^− 1^) < **Delta** (538.1 kcal mol^− 1^) **< Omicron** (749.8 kcal mol^− 1^) using his molecular interaction energy function (Hanai, 2022). This difference from Delta [*ꞵw_min_*[Delta]=-1.22(1)] to Omicron [*ꞵw_min_*[Delta]=-1.35(1)] is measured by FORTE too. Experiments have also shown that the Omicron RBD binds with a 2.4-fold increased affinity to ACE2_wt_ ^(^Cameroni et al., 2021). A recently solved cryo-EM structure of Omicron spike protein–ACE2_wt_ complex confirmed the binding (Mannar et al., 2022). It is worth noting that RBD_Omicron_ was able to maintain its ACE2_wt_ affinity (and overall folded structure) despite a large number of mutations. Although solution pH can affect this complexation, the qualitative comparison is maintained at low and physiological conditions - see Figure 3. Taken together, there is accumulative data for this improved fitness of SARS-CoV-2 RBD for ACE2. The increased RBD_Omicron_-ACE2 attraction can be one of the main important factors contributing to Omicrońs higher transmissibility, reduced incubation time, and ability to establish the infection. This is reinforced by the fact that the viral load is probably not responsible for such high transmissibility (Kozlov, 2022). Comparing Delta and Omicron variants (*βw_min_*[Delta]<*βw_min_*[Omicron] and lower viral load for Omicron in comparison with Delta), it can be suggested that the RBD-ACE2 complexation is directly related to the transmissibility.

**Figure 2:**
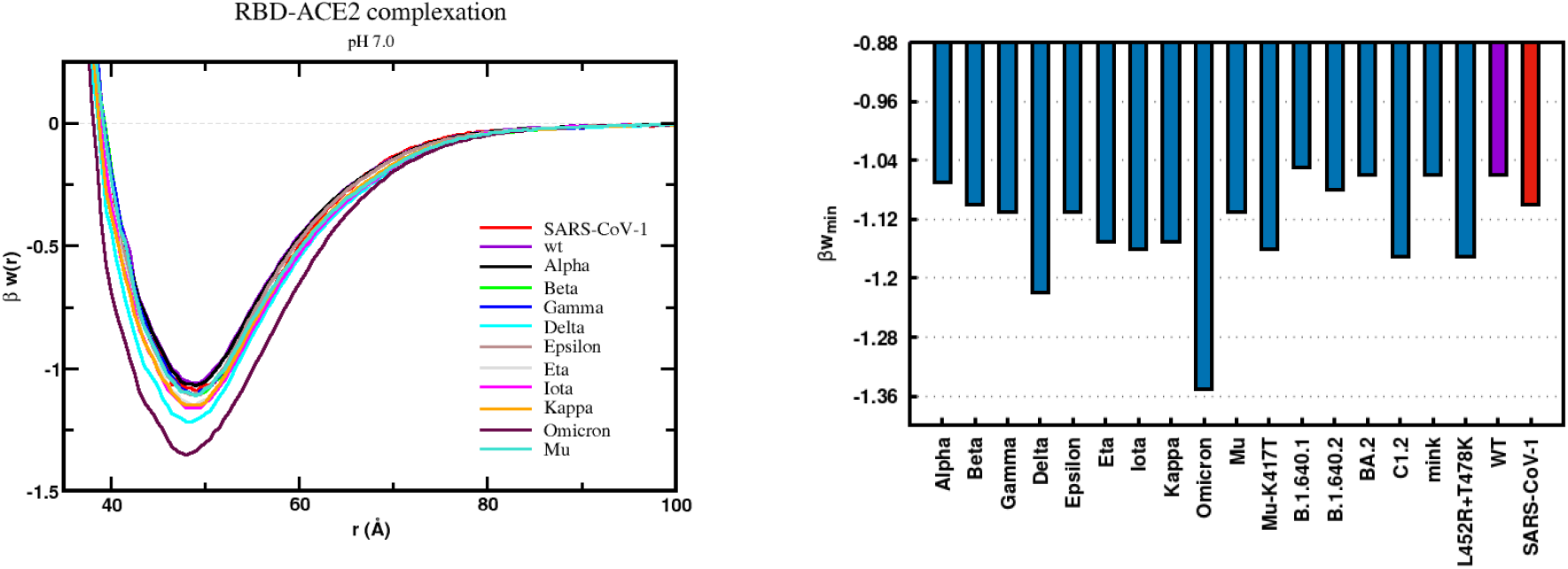
Binding affinities between the RBD from different variants and the cellular receptor ACE2_wt_. (a) *Left panel*: Free energy profiles. The simulated free energy of interactions [βw(r)] between the centers of the RBD proteins from SARS-CoV-1, SARS-CoV-2 (wt), and several variants and the cellular receptor ACE2_wt_ are given at pH 7.0. Salt concentration (NaCl) was fixed at 150 mM. Data for SARS-CoV-1 RBD-ACE2_wt_ and RBD_wt_-ACE2_wt_) was taken from earlier studies (Giron et al., 2020, 2021). Simulations started with the two molecules placed at random orientation and separation distance. (b) *Right panel*: Minima values for each studied system. All *βw_min_* values were taken from the free energy plots (Figure 2a). The maximum estimated error is 0.01K_B_T.

**Figure 3:**
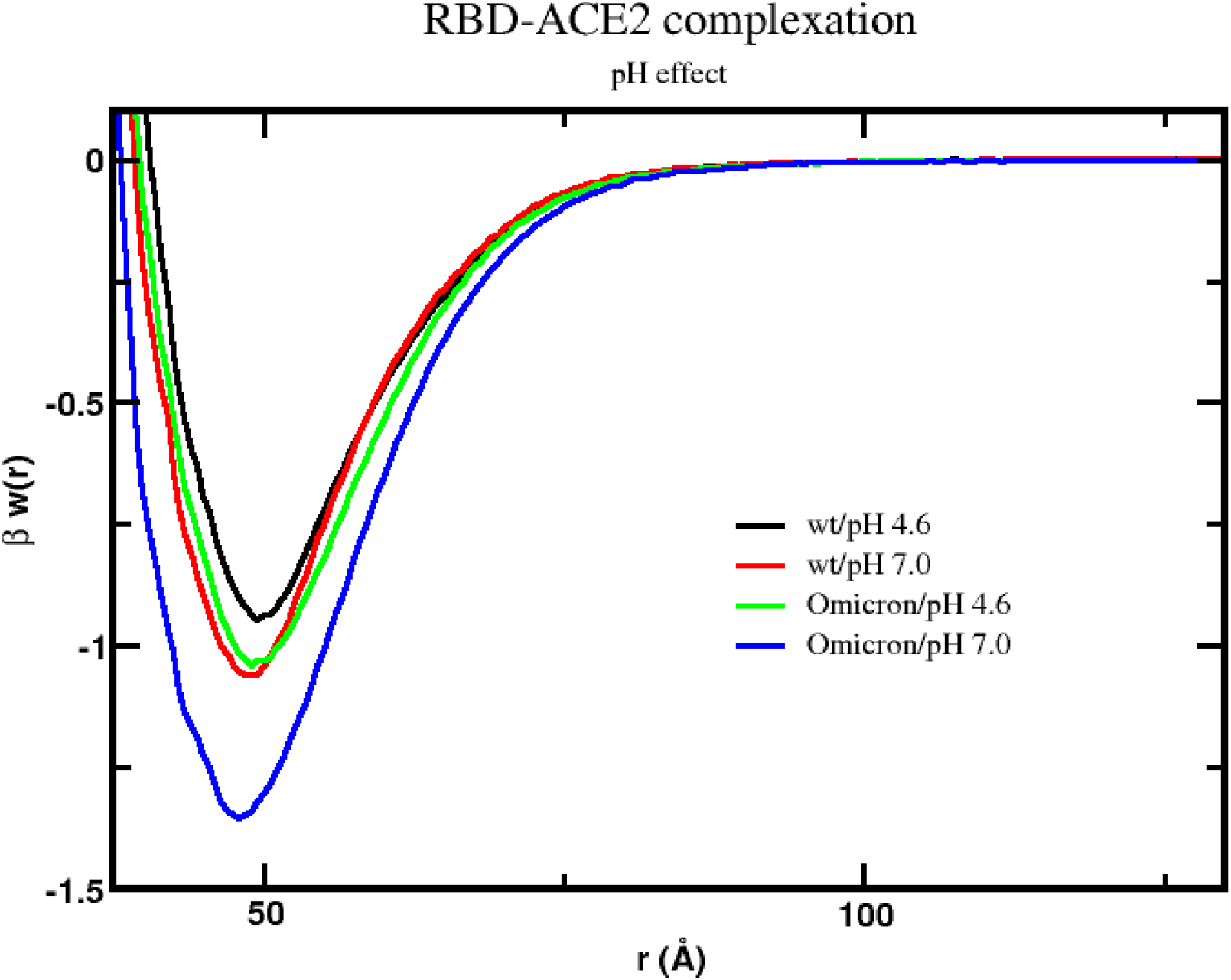
Free energy profiles for the interaction of RBD_wt_ and RBD_Omicron_ proteins with ACE2_wt_. The simulated free energies of interactions [βw(r)] between the centers of mass of the RBD proteins are given at different solution pH conditions. Salt concentration was fixed at 150 mM. Other details as in the previous figure.

A practical need was to map possible synergistic effects among the mutations of the VOCs and VOIs (J. Hu et al., 2022). The presence of more than one mutation does not necessarily imply an increased ACE2_wt_ binding affinity. For instance, both Eta and Iota variants have similar *βw_min_* values for the RBD-ACE2_wt_ complexation [-1.15(1) and −1.16(1), respectively] despite Iota having an additional mutation (N501Y). Conversely, the mutation T478K is key for the Delta variant. Comparing Kappa and Delta variants, the presence of T478K in Delta increased the RBD_Delta_-ACE2_wt_ binding affinity from −1.15(1) (*βw_min_*[Kappa]) to −1.22(1) (*βw_min_*[Delta]).

### 3.2 ACE2 polymorphism

We now turn to the ACE2 polymorphism analyzing if some of them could show a higher binding affinity for the RBD. Although it is not completely established that a higher RBD-ACE2 binding affinity would *directly* imply a higher susceptibility [since other factors can also interfere (e.g. ACE2 can be shed from membranes (Brest et al., 2020)], it can facilitate the virus’s access to the human cell. Therefore, the ACE2 polymorphism has been taught as a possible molecular descriptor to predict the susceptibility for SARS-CoV-2 infection and severity of COVID-19 (Brest et al., 2020). It has some subjectivity with some results indicating an important contribution from this genetic feature (Möhlendick et al., 2021) while others do not show any plausible effect on the risk for infection (Gómez et al., 2020). Despite that, the binding affinities can be measured contributing to the general understanding of the molecular determinants of the virus life cycle. The dbSNP accession number for each ACE2 mutation when known from the literature is provided in parenthesis in this section using the information from the work of Bakhshandeh and colleagues (Bakhshandeh et al., 2021).

In Figure 4, computed free energies of interactions for the principal SARS-CoV-2 RBD variants and ACE2 with common polymorphisms involving the replacement of titratable groups at pH 7 were compiled and displayed as a heatmap-style plot. Qualitatively, Figure 4 reproduces what has already been seen for the ACE2_wt_. The Omicron variant has the strongest binding affinity for any studied ACE2 variant. It is followed by Delta and Kappa. There are ACE2 polymorphisms that increase the RBD-ACE2 binding affinities (e.g. D509Y, G326E, H378R, K26R, and K68E). The mutation K26R (∼0.4% allele frequency (Suryamohan et al., 2021), rs4646116 found in the Italian Cohort (Bakhshandeh et al., 2021)) is known to have an increased binding affinity for RBD_wt_ in experimental assays too (Suryamohan et al., 2021). Among the ACE2 variants that we found with higher RBD-ACE2 binding affinities, H378R (rs142984500, ∼0.01% allele frequency (Suryamohan et al., 2021), common among Europeans (Sayed, 2021)) and K26R were predicted before to increase susceptibility in the case of SARS-CoV-2 wt. This probably means that for these specific ACE2 mutations no other effect plays a significant role to dictate the susceptibility. Conversely, the ACE2 variants D509Y, G326E (rs759579097), and K68E (all with less than 0.001% allele frequency (Suryamohan et al., 2021)) were predicted to be protective ones in the previous analysis for SARS-CoV-2 wt (Suryamohan et al., 2021).

**Figure 4:**
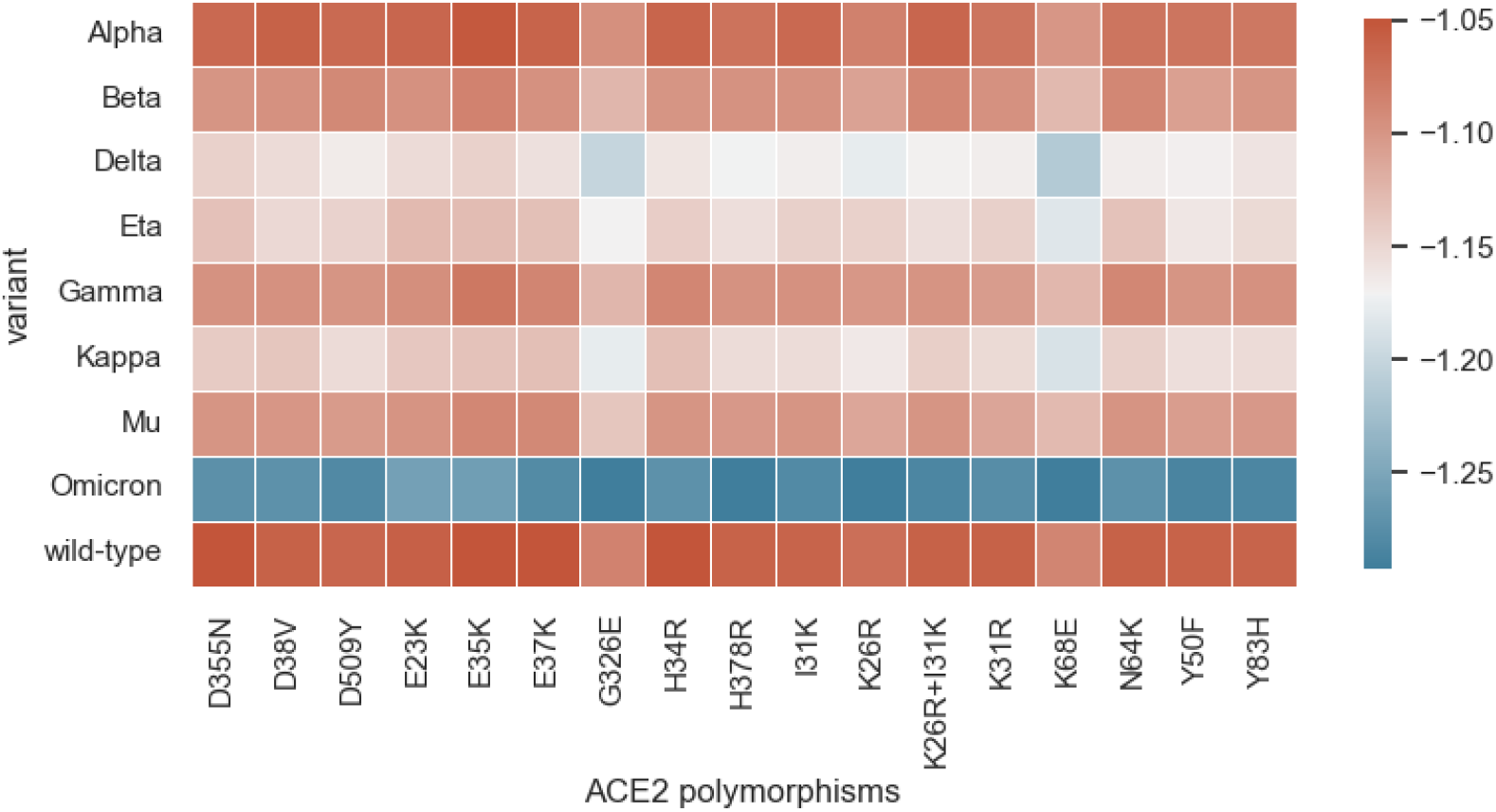
Heatmap with the minima free energy of interactions values measured for the SARS-CoV-2 RBD-ACE2 complexation at pH 7 and 150mM of NaCl for the RBD of the main critical variants and different ACE2 polymorphisms. All values of *βw_min_* are given in K_B_T units. The maximum estimate error is 0.01. See the text for more details.

Both K31R (rs758278442) and E37K (rs146676783, ∼0.004% allele frequency (Suryamohan et al., 2021)) ACE2 polymorphisms were identified as responsible for a decreased RBD_wt_-ACE2 affinity in experimental biochemical assays (Suryamohan et al., 2021). Individuals with these ACE2 polymorphisms would tend to be less susceptible to SARS-CoV-2 infection (in the absence of other factors that could invert it). Previous studies using molecular dynamics in combination with molecular mechanics/Poisson−Boltzmann surface area analysis tested the replacement of K31 and E37 by alanine (K31A and E37A) (Laurini et al., 2020). They confirmed the importance of these two amino acids for the stability of the interface RBD_wt_-ACE2 [the predicted changes in binding free energy were −4.85(0.14) kcal/mol and −2.89(0.15) kcal/mol for RBD_wt_-ACE2(K31A) and RBD_wt_-ACE2(E37A), respectively (Laurini et al., 2020)]. The computed binding affinities in terms of the *βw_min_* values for SARS-CoV-2 RBD wt by FORTE do not show such reduction. The values were −1.06(1), −1.06(1), and −1.05(1) for the binary associations RBD_wt_-ACE2_wt_, RBD_wt_-ACE2_K31R_, and RBD_wt_-ACE2_E37K_, respectively. For Omicron, the reduction is observed: *βw_min_* equal −1.35(1) for RBD_wt_-ACE2_wt_, −1.28(1) for RBD_wt_-ACE2_K31R_, and - 1.29(1) RBD_wt_-ACE2_E37K_.

### 3.3 RBD-mAb free energy of interactions for Omicron and other variants

An issue that has been widely discussed in the literature is the efficacy of the mAbs already used for the wt strain of the virus for the variants, especially for the VOCs (Edara et al., 2021; Hoffmann, Krüger, et al., 2021; L. Liu et al., 2021; Planas et al., 2021; Weinreich et al., 2021). Several new mAbs have also been described in the literature during the last months (Andreano et al., 2021; Du et al., 2021; Jaworski, 2021; Li & Gandhi, 2022). Binders that showed promise in more recently published studies (e.g. COR101, Fab15033, B38, CB6, m336, S2K146, and S309) together with mAbs already known to bind to SARS-CoV-1 were included in the present evaluation of their RBD-mAb complexation for several VOCs and VOIs. Some of them (e.g. m396) were identified in early studies as exhibiting a broad neutralization capacity (Zhu et al., 2007). The list includes mAbs classified (Deshpande et al., 2021) as classes 1 (e.g. B38), 2 (e.g. P2B-2F6), 3 (e.g. REGN10987), and 4 (e.g. CR3022). One anti-N-terminal domain (NTD) mAb (4A8) was also added in our analysis. All mAbs were only tested to bind to the RBDs assumed to be accessible for the complexation (Giron et al., 2020, 2021). Biomolecular interactions with other structural parts of the spike protein were not included in the present analysis. We evaluated how strongly each mAb binds to the virus RBD at a specific physical-chemical condition (pH 7 and 150mM of NaCl) and, consequently, what their *tendencies* for eventual therapeutic potential could be. The neutralization was not quantified in the simulations, and there is no proven correlation between a high binding affinity and a high neutralization (Chi et al., 2020). The binding affinities for some selected illustrative cases of mAbs interacting with the RBD_wt_ are given in terms of βw(r) in Figure 5. Groups of mAbs can be identified from the lowest binding affinity as seen for example for Ab3B4 to the highest affinity observed for S230 that stood out from all the others. However, S230 needs further experimental evaluation to define its risk to enhance virulence since it could contribute to virus-cell fusion (Walls et al., 2019). From the present calculations, it can be seen that S230 indeed mimics ACE2 very well in terms of their free energies of interactions. The binding affinity of S230 to the RBD_wt_ is higher than for the RBD_wt_-ACE2 complexation [*βw_min_*[wt]=-1.06(1) for RBD_wt_-ACE2 complexation and *βw_min_*[wt]=-1.23(1) for RBD_wt_-S230 complexation]. For Omicron, the interaction between the RBD_Omicron_ and S230 is slightly more attractive [*βw_min_*[Omicron]=-1.27(1) for RBD_wt_-S230 complexation] than for SARS-CoV-2 wt while a larger difference is observed for the RBD_wt_-ACE2 complexation [*βw_min_*[Omicron]=-1.35(1) RBD_wt_-ACE2 complexation].

**Figure 5:**
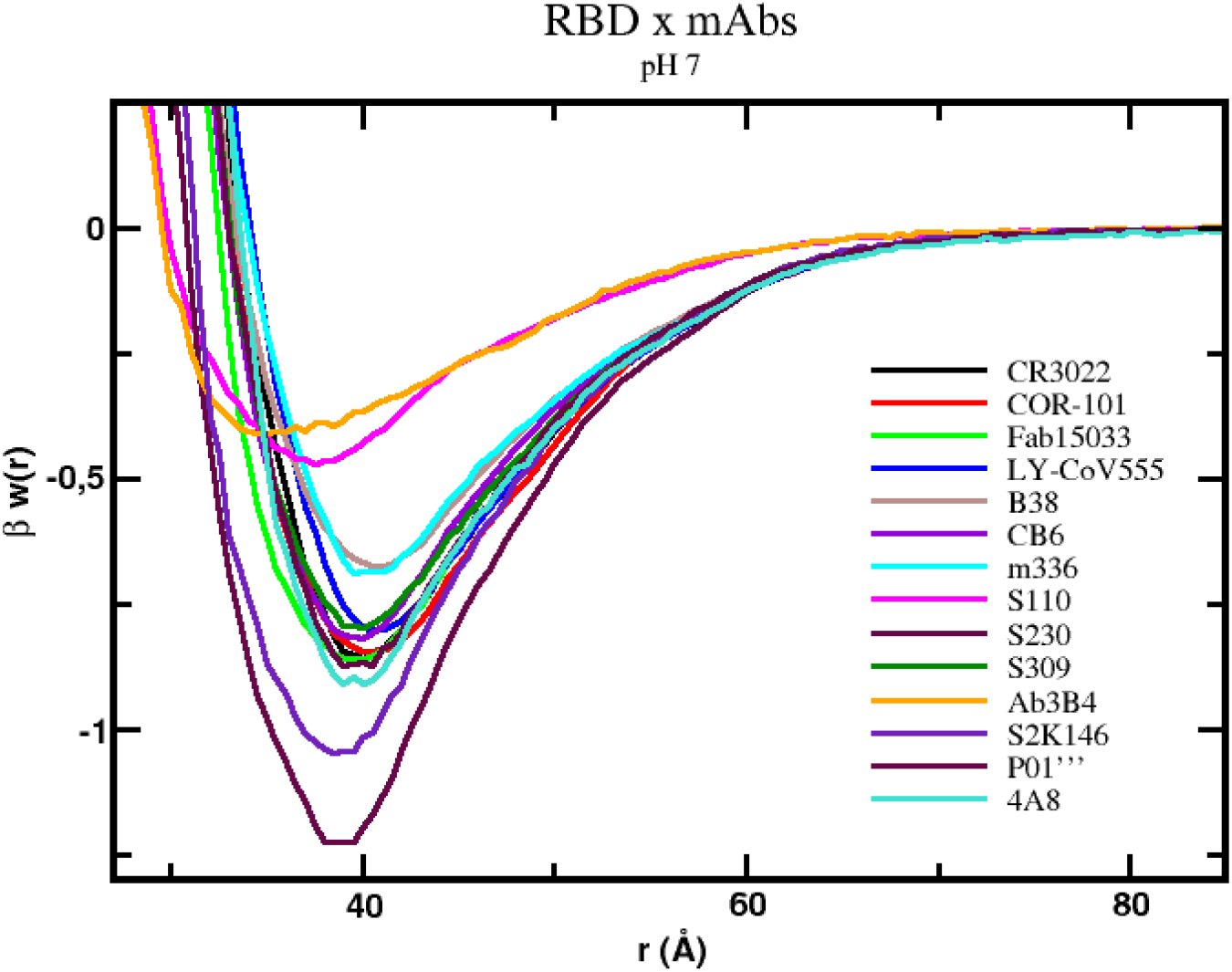
Binding affinities between the RBD_wt_ and some studied mAbs. The simulated free energy of interactions [βw(r)] between the centers of the RBD proteins from SARS-CoV-2 (wt) and the mAbs are given at pH 7.0. Salt concentration (NaCl) was fixed at 150 mM. All other details as in Figure 2.

A full set of data for all studied mAbs is summarized in Figure 6a where a heatmap-style plot displays the values of *βw_min_* for several VOCs and VOIs. In this representation, as dark is the blue as higher is the RBD-mAb affinity while as dark is the red as lower is the binding affinity. Besides S230, S2K146 is another mAb that also stood out from the others in terms of its binding affinity. This suggests S2K146 as a potentially effective alternative to bind and eventually neutralize the virus interaction with the cellular receptor and, consequently, to be used as treatment or prophylaxis. S2K146 has the second-highest RBD binding affinity in comparison with all studied mAb under the investigated physical-chemical conditions. Moreover, S2K146 binds with such high affinity to all investigated RBDs including SARS-CoV-1 [*βw_min_*[wt]=-1.06(1)] and Omicron [*βw_min_*[Omicron]=-1.10(1)]. S2K146 can also mimic ACE2. For instance, the value for the free energy of interaction for the RBD_wt_-S2K146 binding [*βw_min_*[wt]=-1.05(1)] is equivalent to the one for the RBD_wt_-ACE2_wt_ complexation [*βw_min_*[wt]=-1.05(1)] while the RBD_Omicront_-ACE2_wt_ complexation [*βw_min_*[Omicron]=-1.35(1)] is more attractive than the RBD_Omicron_-S2K146 association [*βw_min_*[Omicron]=-1.10(1)]. Cameroni and co-authors reported experimental values of IC_50_ equals 14.2 and 12.6 ng/ml for the wt and the Omicron variant, respectively (Cameroni et al., 2021). Such broadly binding capacity has also been experimentally shown both for the wild-type, as well as the Alpha, Beta, Delta, Epsilon, and Kappa variants, (Park et al., 2022) and Omicron (Cameroni et al., 2021). The neutralization power of S2K146 was experimentally measured for the wt and some variants (IC_50_ equals 10 ng/ml for the wt, 9 ng/ml for Alpha, 9 ng/ml for Delta, and 30 ng/ml for Kappa (Park et al., 2022)) reproducing similar tendencies to what was computed here by FORTE for the RBD-S2K146 binding affinities [*βw_min_*[wt]=-1.05(1); *βw_min_*[Alpha]=-1.07(1); *βw_min_*[Delta]=-1.05(1); *βw_min_*[Kappa]=-1.04(1)]. This is promising for clinical trials. In fact, *in vivo* tests with hamsters infected with SARS-CoV-2 have also shown that S2K146 is capable of greatly reducing or eliminating viral replication (Park et al., 2022).

**Figure 6:**
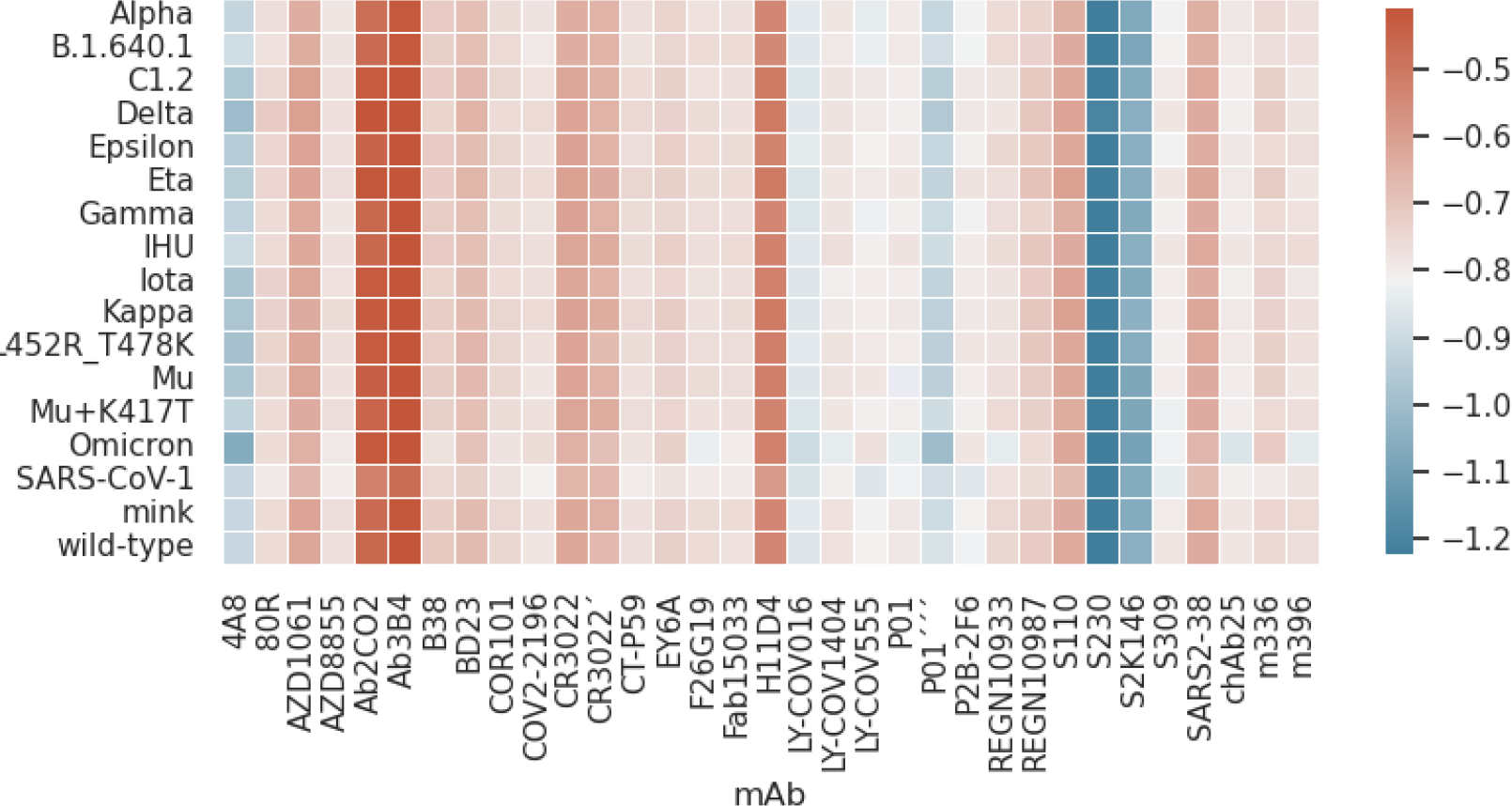
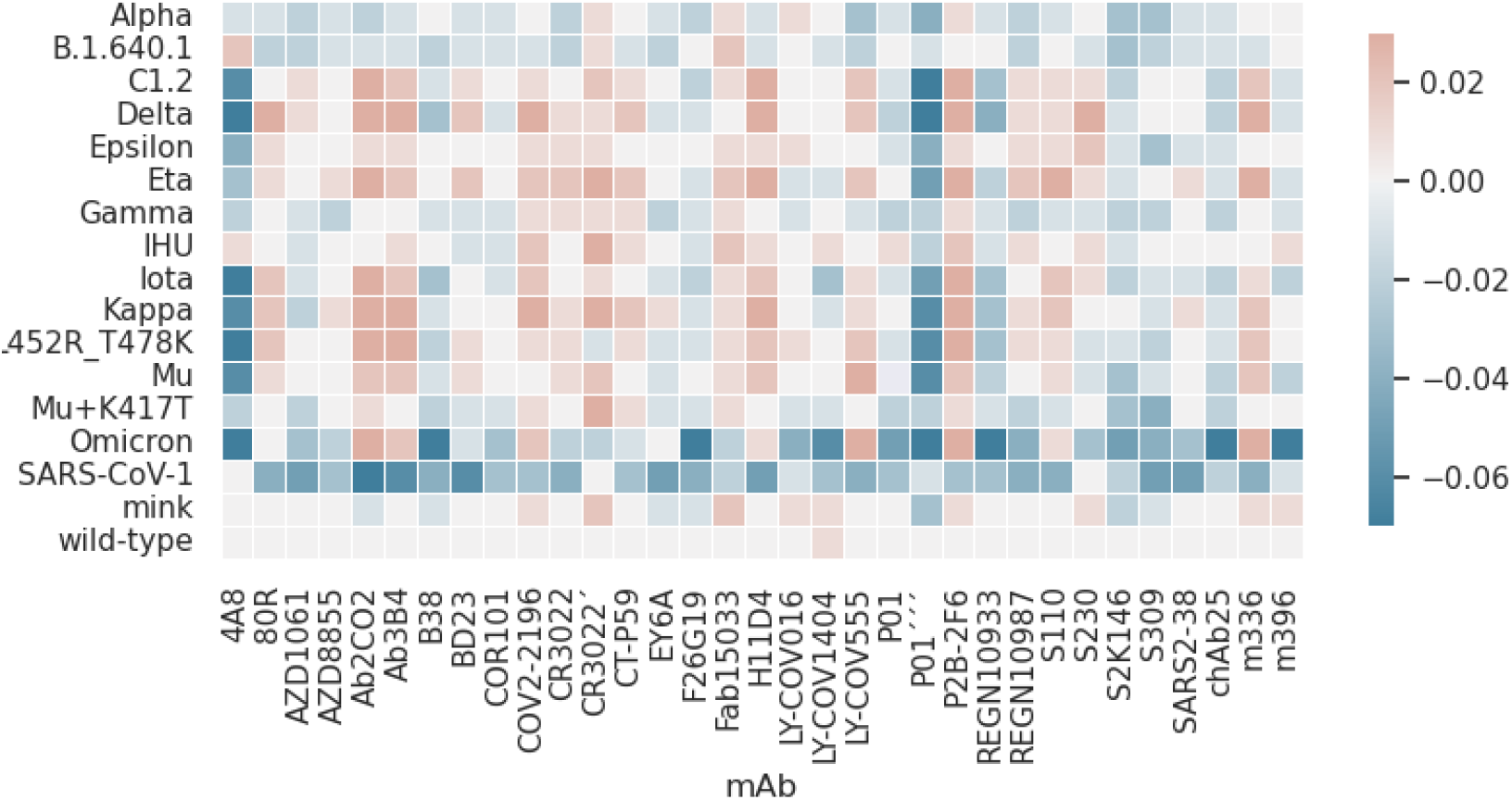
Heatmap with the minima free energy of interactions values measured for the SARS-CoV-1 and 2 RBDs-mAbs complexations at pH 7 and 150mM of NaCl. Data for all studied systems. Other details are as in Figure 4. (**a**) *Top panel*: absolute values of *βw_min_*. (**b**) *Bottom panel*: the differences between *βw_min_*[variant] and *βw_min_*[wt] (ΔA=*βw_min_*[variant] and *βw_min_*[wt]).

Ranking the RBD-mAbs complexation from the top to the bottom (using their *βw_min_* values), S2K146 [*βw_min_*[wt]=-1.05(1)] is followed by the anti-NTD 4A8 [*βw_min_*[wt]=-0.91(1)] and the engineered anti-RBD P01’’’ [*βw_min_*[wt]=-0.87(1)] in terms of their binding affinities. Yet, this predicted result for 4A8 does not totally agree with the experimental findings. 4A8 is known to bind to the NTD with high affinity (K_D_ = 92.7 nM) and a weaker affinity for the RBD (EC_50_ > 10,000 ng/ml in an ELISA experiment) (Chi et al., 2020). Computer simulations confirmed the 4A8-NTD complexation (K_D_ = 9.1 nM) but did not test before the possibility it could also bind to the RBD (Nguyen et al., 2021). On the basis of the present calculations, the 4A8 binding to NTD might not be unique since the anti-NTD 4A8 mAb could also bind to the RBD from the VOCs and VOIs with high affinity at pH 7 and 150mM. Such discrepancy could be due to the different experimental physical-chemical conditions or the presence of any other molecular mechanisms not included in the simulations (e.g. the up and down conformational state changes, repulsion from another structural region of the spike protein absent in the model, etc.). It was speculated that the neutralization mechanism of 4A8 was through restraining the spike protein at the down conformational state (Chi et al., 2020). We could add that the 4A8 binding to the RBD could also be part of the explanation based on the present data assuming the spike protein can be at the up configurational state. This hypothesis can be further experimentally tested by performing neutralization assays with Omicron. For Omicron, a better neutralizing effect is expected since the attraction was increased for RBD_Omicron_ in comparison to the wt. If the antibody-dependent enhancement of infectivity (typically observed for anti-NTD mAbs like 4A8 (Hachim et al., 2021; Y. Liu et al., 2021)) would be absent for 4A8, it could simultaneously target two different regions of the spike protein which might increase its neutralization quality by both reducing the interaction with ACE2 and allowing the virus particle to be disposed of by a specialized immune cell.

Figure 6b is similar to 6a but highlights the effect of the mutations from each variant on the binding affinities for a given mAb. The difference between *βw_min_*[variant] and *βw_min_*[wt] (ΔA=*βw_min_*[variant] and *βw_min_*[wt]) is plotted instead of the absolute *βw_min_* values given in Figure 6a. While Figure 5 illustrates the diversity of free energy profiles, Figures 6a and 6b allow a comparison of all studied mAbs among different VOCs and VOIs. It can be noted that the affinity is slightly affected for each variant. For a given mAb, the interaction with the RBD of each variant has its own features. This also helps to understand how is the immune response produced by vaccines that each one should drive the human body to produce specific Abs after the immunization. From the data given in Figure 6a and taking into account the estimated errors, the studied mAbs can be ranked in terms of their binding affinities for the RBD_wt_ as group 1 (Ab3b4) < group 2 (Ab2CO2) < group 3 (H11D4, and S110) < group 4 (ADZ1061, CR3022, and SARS2-38) < group 5 (CR3022’, and BD23) < group 6 (B38, REGN10987, and EY6A) < group 7 (COR-101, REGN10933, m336, F26G19, 80R, m396, CT-P59, ADZ8855, Fab15033, LY-CoV-1404, COV2-2196, chAb25, S309, and P01) < group 8 (LY-CoV555, and P2B-2F6) < group 9 (LY-CoV016, and P01’’’) < group 10 (4A8) < group 11 (S2K146) < group 12 (S230). Omicron behaves similarly to the wildtype for groups 1-5. From group 5 on, the rank for Omicron is group 6 (m336, and EY6A) < group 7 (COR-101, REGN10987, F26G19, 80R, CT-P59, ADZ8855, Fab15033, COV2-2196, LY-CoV555, and B38) < group 8 (S309, F26G19, P01, LY-CoV1404, REGN10933, m396, and chAb25) < group 9 (LY-CoV016) < group 10 (P01’’’) < group 11 (4A8) < group 12 (S2K146) < group 13 (S230). For all VOCs and VOIs, LY-CoV016, P01’’’, 4A8, S2K146, and S230 are the mAbs with the highest RBD binding affinities and cross-reactivity. For the sake of comparison, the *βw_min_* values of these top 5 mAbs are shown in Figure 7 for wt, Delta, and Omicron variants.

**Figure 7:**
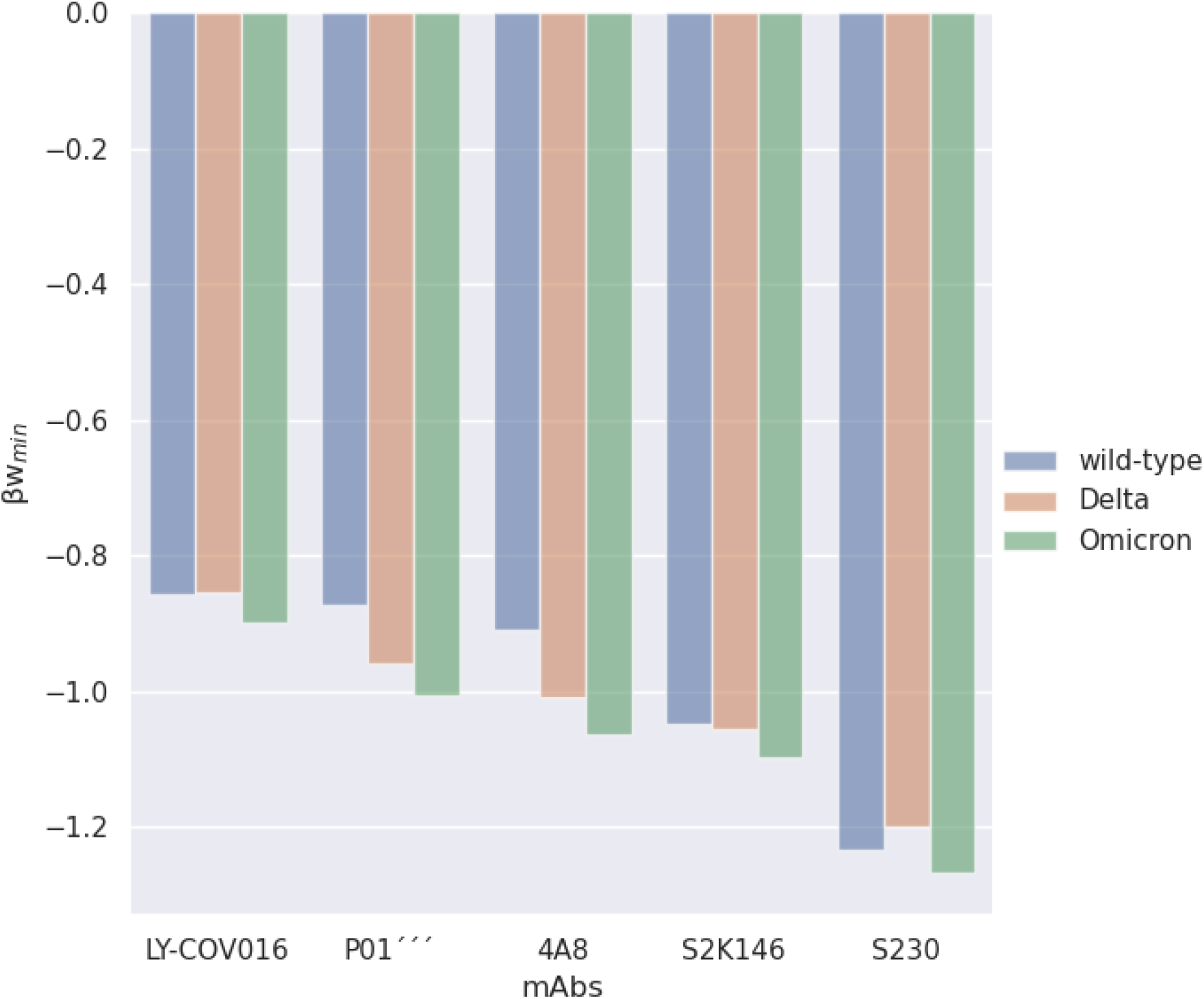
Binding affinities between the RBD from different variants and the mAbs with the highest binding affinities from all the antibody landscapes. Data from simulations at pH 7.0 and 150mM of NaCl.

Omicron and Delta variants tend to share the same behavior in terms of RBD-mAbs affinities, but there are exceptions (see Figures 5 and 6). For instance, the mutations in Omicron favor the RBD_Omicron_-S230 interaction, while the opposite is seen for RBD_Delta_-S230. In other cases, a mAb can be more attracted for RBD_Omicron_ (ΔA=-0.04 for the RBD_Omicron_-LY-CoV016 complexation) while the mutations in Delta do not show a change (ΔA=0.0 for the RBD_Delta_-LY-CoV016 complexation). For most of the studied mAbs, the mutations in Omicron can increase the RBD_Omicron_-mAb affinity except for Ab2CO2, Ab3B4, COV2-2196, LY-CoV555, P2B-2F6, and m336). Other factors besides the RBD binding affinity (not included here) might affect the *in vitro/in vivo* viral neutralization of some mAbs. Although no reduced effect was found in the computed free energies by FORTE involved in the RBD-mAb complexation, it was reported that the neutralization potency of LY-CoV016, REGN10933, REGN10987, and AZD1061 have a great reduction in their neutralization potency by Omicron (Cao et al., 2021). For other systems (LY-CoV555 and COV2-2196/AZD8895), the decrease in the RBD_Omicron_-mAb binding affinity follows the reduction in the neutralization by Omicron (Cao et al., 2021).

S309 appears in group 8 closer to the best candidates with *βw_min_* values −0.79(1) and −0.82(1) for the complexations RBD_wt_-S309 and RBD_Omicron_-S309, respectively. Theoretical predictions by Yang and colleagues indicated that this mAb could still neutralize the Omicron variant (Q. Yang et al., 2021). Our calculations (see Figure 6a) showed that the RBD_Omicron_-S309 complexation is slightly more attractive at the studied physical-chemical conditions than the RBD_wt_-S309 complexation which corroborates this previous study. Comparing the present data from the simulations for S309 and the top 5 mAbs listed above (LY-CoV016, P01’’’, 4A8, S2K146, and S230 - see Figure 7) suggest a more successful use of them in clinical trials than Sotrovimab, the mAb already authorized by FDA (Gandhi et al., 2022). Though S230 has the highest affinity among all, it might be the only exception due to its risk of antibody enhancement (not evaluated in the present work) (Denner, 2020; Starr et al., 2020).

Structural comparisons among some putative binding modes for each of the five best mAbs as illustrated in Figure 8 are suggestive for their combined practical use. Figure 8 suggests that **i**) S2K146 could behave similarly to S230 due to its putative binding region almost overlapping the one observed for S230 (see the red and green molecule in the top of Figure 8), and **ii**) different structural parts of the spike protein can be simultaneously targeted. Therefore, it would be a promising route to further explore them in experimental assays and clinical trials. On the basis of previous data for Influenza and HIV and recent results already available for Omicron, cocktails of mAbs seem a safer bet than monotherapies (Rockett et al., 2021). S2K146 (providing S2K146 does not trigger fusogenic conformational changes as possible for S230) together with P01’’’ and LY-CoV-016 seem to be good candidates for such approach due to their strong binding affinities.

**Figure 8:**
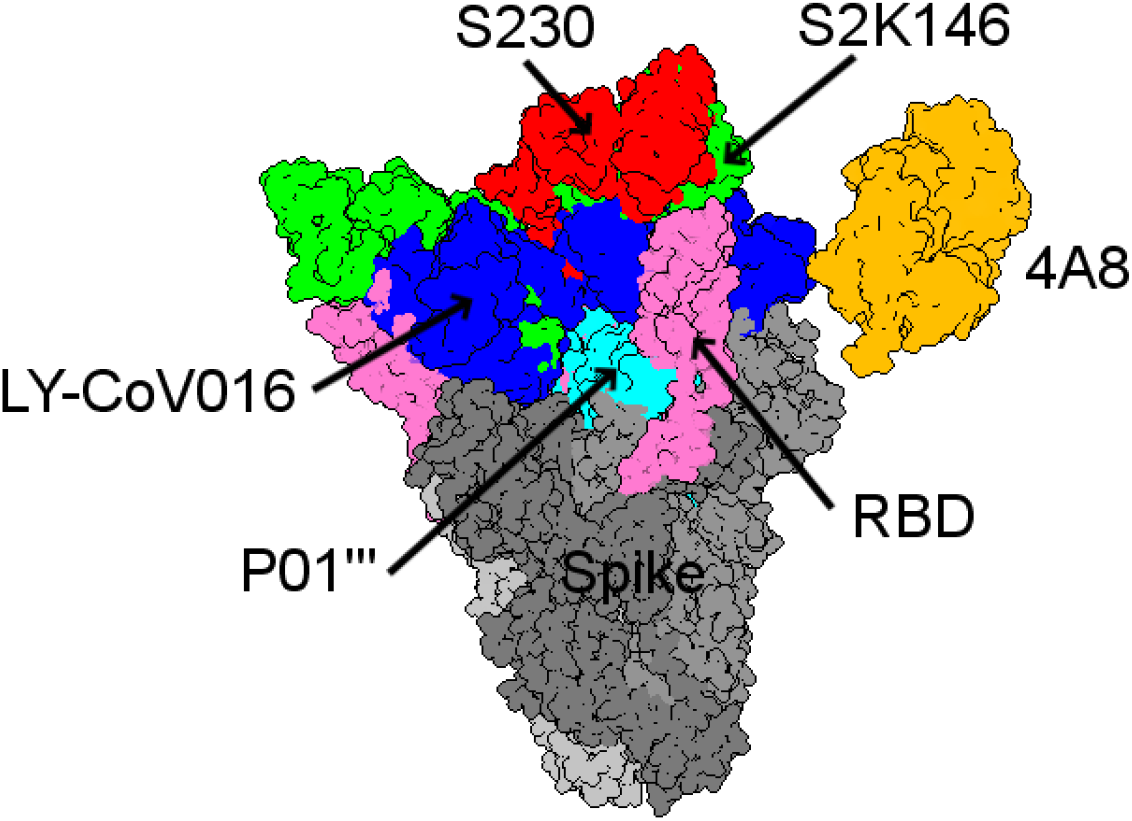
Structural comparison between the five best mAbs with the highest binding affinities from all the antibody landscapes. The spike protein as given by PDB id 7TAT is used as a general template for all putative mAbs [shown in red (S230), green (S2K146), orange (4A8), cyan (P01’’’) and blue (LY-CoV016)]. The RBD region of the spike homotrimer was colored in pink.

## 4. Conclusions

The solution pH together with temperature and pressure are important thermodynamic variables in experimental studies and used by our cells in their biochemical functions. In our computer simulations, we can change the pH (used as an input parameter) and the charges of titratable groups following their protonation and deprotonation. The pH-dependent valences of the titratable groups are controlled by the fast proton titration scheme (FTPS) (Barroso da Silva et al., 2017). In our simulation model also the hydrophobic effect, van der Waals interactions, and excluded volume repulsion, through an LJ term between the residues are included. We have previously applied this methodology successfully in several biomolecular systems (Barroso da Silva et al., 2018; Delboni & Barroso da Silva, 2016; Mendonça et al., 2019; Persson et al., 2010) and to investigate the flaviviruses (Poveda-Cuevas et al., 2018, 2020) and coronavirus (Giron et al., 2020, 2021; Neamtu et al., 2022).

After recent updates, the method is named FORTE (**F**ast c**O**arse-grained p**R**otein-pro**T**ein mod**E**l) (Neamtu et al., 2022). Unlike docking, the FORTE approach provides derivatives of the free energy, e.g., the interaction free energy as a function of separation distances but not limited to a specific separation/orientation between the macromolecules. The model considers each amino acid as a single charged Lennard-Jones sphere of varied radius R_i_ proportional to the effective size of the residue with valence z_i_. The initial coordinates are given by its three-dimensional structures allowing non-specific hydrophobic effects. Electrostatic interactions are properly treated as they are of great importance in promoting interactions between macromolecules and even more critical for virus proteins.

We have applied here the FORTE coarse-grained model and simulation tool originally developed for protein-protein interaction studies at constant pH conditions to study the SARS-CoV-2 RBD interactions with cell receptors and mAbs. We focused on the binding affinities as important molecular determinants of the virus life cycle. At the same time, ACE2 polymorphism is a possible molecular descriptor to predict the susceptibility for SARS-CoV-2 infection and the severity of COVID-19.

The main goal in our work has been to compare Omicron with other strains of how its RBD binding affinities in the complexation with ACE2 polymorphs and the mAbs. To our best knowledge, no previous studies have so far investigated and compared the RBD-ACE2 and RBD-mAbs binding affinities for such many cases at the same conditions using simultaneously a common *in silico* strategy for all of them. Totally 33 binders and groups of mAbs are identified for our study. All the mAbs showing strong binding capacity can also bind to the RBD from SARS-CoV-1, SARS-CoV-2 wt, including all now studied variants.

In our study, we assume the spike protein found in the “open” configuration and therefore highly accessible to ACE2 and/or the other binders. Some conformational adjustments might be needed in the spike homotrimer for the *in vivo* complexation RBD-ACE2 without any steric clashes. We find that the sequence dependence on the structure may have some potential role in introducing modifications in the interactions while the overall coiling is still kept preserved. Conformational changes have a minor effect on the binding free energies in particular for medium and long-range separation distances.

S2K146 binds strongly and potentially nullifies the virus interaction with the cellular receptor and, could be used both preventively and as treatment. In our study S2K146 had the second-highest RBD binding affinity among all studied mAbs, binding with high affinity to all investigated RBDs as well as SARS-CoV-1. Omicron has the strongest binding affinity for any studied ACE2 variant followed by Delta and Kappa. We find that Omicron and Delta variants share mutually similar behavior concerning the RBD-mAbs affinities with some exceptions.

The five best mAbs candidates from our calculations with the highest binding affinities are S230, S2K146, P01’’’, 4A8, and LY-CoV-016. S2K146 and S230 have similar binding affinities due to their nearly overlapping binding region but they are still able to target different structural parts of the spike protein. Rather than applying them separately, S2K146 together with P01’’’ and LY-CoV-016 could provide an efficient cocktail based on their strong individual binding affinities. 4A8 is a good candidate as it seems to target two different regions of the spike protein simultaneously thereby increasing its neutralization power.

## Acknowledgments

This work has been supported in part by the “Fundação de Amparo à Pesquisa do Estado de São Paulo” [Fapesp 2020/07158-2 (F.L.B.d.S.)] and the Conselho Nacional de Desenvolvimento Científico e Tecnológico (CNPq) [CNPq 305393/2020-0 (FLBdS)]. F.L.B.d.S. is also deeply thankful for resources provided by the Swedish National Infrastructure for Computing (SNIC) at NSC and PDC. A. Laaksonen acknowledges the Swedish Research Council for financial support, and partial support from a grant from the Ministry of Research and Innovation of Romania (CNCS -UEFISCDI, project number PN-III-P4-ID-PCCF-2016-0050, within PNCDI III). The authors also gratefully acknowledge the computing time granted by the John von Neumann Institute for Computing (NIC) and provided on the supercomputer JURECA at Jülich Supercomputing Centre (JSC).

## Acknowledgments

The financial support by the Fundação de Amparo à Pesquisa do Estado de São Paulo [Fapesp 2020/07158-2 (FLBdS)], the Conselho Nacional de Desenvolvimento Científico e Tecnológico [CNPq 305393/2020-0 (FLBdS) and PIBIC/CNPq 2020-1732 (CCG)], the Swedish National Infrastructure for Computing (SNIC) at NSC, the Swedish Research Council, and from the Ministry of Research, and Innovation of Romania (CNCS-UEFISCDI, project number PN-III-P4-ID-PCCF-2016-0050, within PNCDI III) are much appreciated. The authors also gratefully acknowledge the computing time granted by the John von Neumann Institute for Computing (NIC) and provided on the supercomputer JURECA at Jülich Supercomputing Centre (JSC).

